# Genome editing of the flavanone 3-hydroxylase gene produces a green *Perilla* line with elevated luteolin accumulation

**DOI:** 10.64898/2025.12.28.695628

**Authors:** Shuji Matsushita, Michiharu Nakano, Suguru Chokyuu, Masaki Kurao, Ayane Fujita, Junko Kimura, Chinatsu Nagata, Takeshi Ishikawa, Keita Tamura, Hidemasa Bono

## Abstract

*Perilla frutescens* var. *crispa* is widely cultivated in East Asia and produces a diverse range of secondary metabolites with nutritional and medicinal value. Based on anthocyanin content, perilla is classified into red and green chemotypes. The anthocyanin biosynthetic pathway is highly conserved among plants, and the flavanone 3-hydroxylase gene (*F3H*) in red perilla is considered essential for this pathway. Here, we used CRISPR–Cas9 genome editing to generate novel *Perilla* lines with altered metabolite profiles. In four edited T₀ plants, multiple biallelic indels, including a −3 bp in-frame deletion, were introduced at the target site. In the T₁ generation, these mutations and the transgene segregated in a Mendelian manner, resulting in recovery of null-segregant plants. Metabolite analysis revealed strong suppression of anthocyanin biosynthesis and a significant increase in luteolin accumulation in edited lines compared with the wild type. Transcriptome analysis further confirmed the loss of *F3H* function, demonstrating a metabolic shift of the flavonoid pathway toward luteolin production. This study provides the first functional validation of *F3H* in perilla and highlights its potential application in breeding cultivars with controlled bioactive compound contents. Our findings also demonstrate practical strategies for implementing genome editing in non-model crops and developing high-value functional cultivars.

**Practitioner Points:**

- Editing a single flavonoid biosynthetic gene (*F3H*) effectively alters pigment composition in perilla leaves.
- Increased accumulation of luteolin-related compounds suggests potential for enhancing functional metabolites without complex pathway engineering.
- Genome editing offers a feasible tool for perilla improvement in breeding and functional food development.

## Introduction

The genus *Perilla* L. (Lamiaceae) comprises annual, self-pollinating plants that are widely distributed and cultivated in East Asia, including in China, Korea, and Japan (Nitta et al., 2005). Several varieties are recognized depending on their use and morphology. *Perilla frutescens* var. *frutescens* is mainly grown as an oilseed crop (*egoma*) for the purpose of extracting oil from its seeds, and its leaves are consumed in Korea. In contrast, *P. frutescens* var. *crispa* is widely cultivated as a culinary herb and is divided into two types, namely, red *(aka-shiso*) and green (*ao-jiso*), according to leaf and stem pigmentation. In Japan, red perilla is commonly used to color and flavor pickled plums, with anthocyanins responsible for the red pigmentation (Nitta et al., 2003), and its leaves are often dried and ground into a rice-seasoning (*furikake*) powder. In contrast, green perilla is primarily used fresh as a garnish for sashimi, providing both flavor and visual appeal, and is regarded as indispensable in Japanese cuisine.

*Perilla* contains over 400 identified bioactive compounds, which give it considerable value as a functional food and medicinal resource (Hou et al., 2022). In traditional Chinese medicine, dried perilla leaves are widely used to treat gastrointestinal disorders, colds, and anxiety (Ahmed, 2019). Notable bioactive constituents include perillaldehyde, a monoterpenoid with reported antidepressant, anticancer, and antibacterial effects (Uemura et al., 2018); anthocyanins, flavonoids with strong antioxidant activity; luteolin, which exhibits both antioxidant and anti-inflammatory effects (Ueda et al., 2002; Jeon et al., 2014; Kim & Lee, 2019a); and rosmarinic acid, particularly abundant in green perilla and known for its potent anti-inflammatory and antioxidant functions (Deguchi & Ito, 2020). Overall, these compounds contribute to health maintenance and disease prevention, making the regulation of their accumulation directly relevant to breeding high-value cultivars and functional food resources.

*P. frutescens* is an allotetraploid species (2n = 4x = 40), originating from natural hybridization between the diploid species *P. citriodora* (2n = 20) and a presently identified unidentified wild relative (Ito & Honda, 1999; Ito & Honda, 2007; Zhang et al., 2021). This genomically complex structure contributes to the production of diverse secondary metabolites, while posing substantial challenges for genetic analysis. In contrast, advances in molecular biological approaches have accelerated the research in *Perilla*, leading to the elucidation of secondary metabolic pathways and the cloning of related genes (Gong et al., 1997; Gong et al., 1999a; Saito & Yamazaki, 2002). With the rapid development in sequencing technologies, comparative transcriptome analyses of the red and green perilla have been conducted (Fukushima et al., 2015), and more recently, quantitative trait locus analyses based on segregating populations have been reported (Kinoshita et al., 2023; Xie et al., 2025). Furthermore, our group has recently achieved a high-quality reference genome assembly for red perilla (Tamura et al., 2023), increasing the genetic and genomic resources available for breeding.

In recent years, genome editing technologies, particularly CRISPR–Cas9, have been increasingly used to enhance secondary metabolite production in medicinal and functional crops (Das et al., 2024; Rynjah et al., 2025; Sagharyan et al., 2025). In perilla, editing of the *HD3a* gene, a key regulator of photoperiodic flowering, successfully delayed flowering and increased leaf yield (Yun et al., 2023). However, genome editing aimed at modifying the accumulation of bioactive metabolites has not yet been reported (Karthik et al., 2024). In perilla, the *F3H* gene encodes flavanone 3-hydroxylase, which catalyzes the conversion of naringenin to dihydrokaempferol, a key step in anthocyanin biosynthesis (Saito & Yamazaki, 2002). Loss of *F3H* function has been shown to reduce or abolish anthocyanin accumulation in multiple plant species, including strawberry and mulberry, through impaired flavonoid biosynthesis (Dai et al., 2022; Xu et al., 2023). Based on these observations, loss of *F3H* function in red perilla is expected to block anthocyanin accumulation, resulting in a green perilla-like phenotype, and may also alter flavonoid flux, potentially affecting the accumulation of other functional metabolites such as luteolin.

In this study, we targeted the *F3H* gene in red perilla using the CRISPR–Cas9 system to generate novel lines with modified metabolite composition through the suppression of anthocyanin biosynthesis and altered luteolin content. Our findings provide a new strategy for improving the functional properties of perilla and offer an important case study for the application of genome editing in polyploid crop species.

## Results

### Generation of genome-edited perilla plants

Hypocotyl explants (n = 239) of *Perilla frutescens* ‘Hoko-3’ were inoculated with *Agrobacterium tumefaciens* strain EHA105 carrying the genome-editing vector. Following callus induction, shoot regeneration under tissue culture conditions, and kanamycin-based selection, four green regenerants were obtained. PCR analysis confirmed vector integration in all regenerated plants, and Sanger sequencing verified the presence of targeted mutations. However, due to the allotetraploid nature of *P. frutescens*, precise characterization of all allelic mutations in each plant was not feasible (Fig. 1B).

**Fig. 1.**
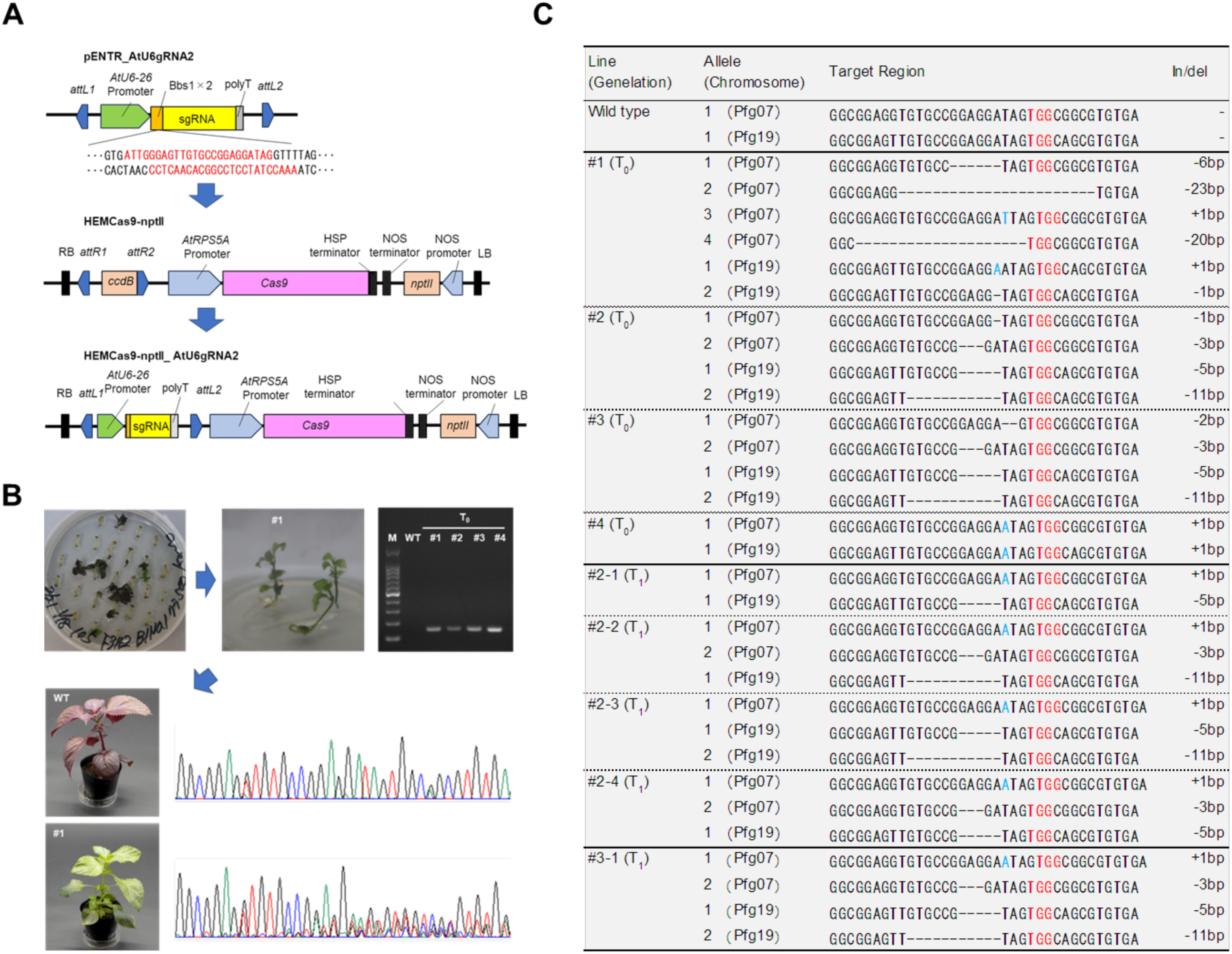
Generation of genome-edited plants. (A) gRNA oligos were inserted into pENTR_AtU6gRNA2, and the gRNA cassette was transferred into HEMCas9-nptII by Gateway recombination to generate the final genome editing vector. (B) Generation of T₀ transgenic plants. Shoots were regenerated from hypocotyl explants after *Agrobacterium* infection via callus formation and were excised for root induction. Individuals that developed roots were acclimated after transformation was confirmed by PCR, followed by Sanger sequencing. (C) Confirmation of the edited region. Target region of genome editing was finally validated by amplicon sequencing.

Amplicon sequencing further characterized the edited loci, which were assigned to linkage groups Pfg07 and Pfg19 based on the reference genome assembly (*Pfru_yukari_1.0*; Tamura, 2023), suggesting that the two loci are likely located on different chromosomes and may segregate independently. In the T₀ generation, mutations ranging from –23 bp deletions to +1 bp insertions were detected (Fig. 1C, Table S1). Line #1 carried four distinct mutations at Pfg07 and two at Pfg19, indicative of a chimeric genome. Lines #2 and #3 exhibited identical –3 bp deletions at Pfg07, suggesting potential partial restoration of gene function. In line #4, +1 bp insertions were observed at both Pfg07 and Pfg19 alleles, resulting in complete biallelic mutations.

Self-pollination of lines #1–#4 generated T₁ seeds. Amplicon sequencing of progeny from lines #2–#4 revealed stable inheritance of edited alleles. In line #2, six progeny were analyzed: the –3 bp deletion at Pfg07 and the –5 bp and –11 bp deletions at Pfg19 were stably transmitted, whereas the – 1 bp deletion at Pfg07 in the T₀ generation was replaced by a +1 bp insertion. In line #3, all 14 analyzed progeny consistently inherited the –3 bp mutation at Pfg07, confirming segregation from the T₀ generation. In line #4, the +1 bp insertions at both Pfg07 and Pfg19 were fixed as homozygous alleles in the progeny.

### Metabolite profiling of genome-edited perilla plants

T₁ individuals from lines #2 and #3 were clonally propagated in vitro and subsequently hydroponically cultivated in an isolated greenhouse. A single hydroponic bed was used to grow both the wild type (WT, non-edited ‘Hoko-3’) and a green *P. frutescens* cultivar as controls. This arrangement ensured a uniform genetic background and consistent growth conditions, enabling precise comparative analyses. Anthocyanin, polyphenol, and other metabolite content of the leaves was measured.

Genome-edited plants with disrupted *F3H* function predominantly exhibited green leaves, in contrast to the strong red pigmentation along leaf veins observed in WT (Fig. 2). Lines carrying the −3 bp deletion showed faint purple coloration along the veins. HPLC analysis of leaves harvested in June and September confirmed that these morphological differences were reflected in the metabolite profiles. Across both sampling dates, WT accumulated 2,814–3,628 µg/g FW of anthocyanins, whereas edited lines showed drastic reductions in anthocyanin content. Complete loss-of-function lines (#2-1 and #2-3) accumulated only 15–18 µg/g FW, whereas the −3 bp deletion lines (#2-2, #2-4, #3-1) retained slightly higher levels (approximately 56–126 µg/g FW), indicating partial *F3H* activity. Overall, most edited plants had anthocyanin contents below 100 µg/g FW, consistent with their reduced or absent purple pigmentation (Fig. S2).

**Fig. 2.**
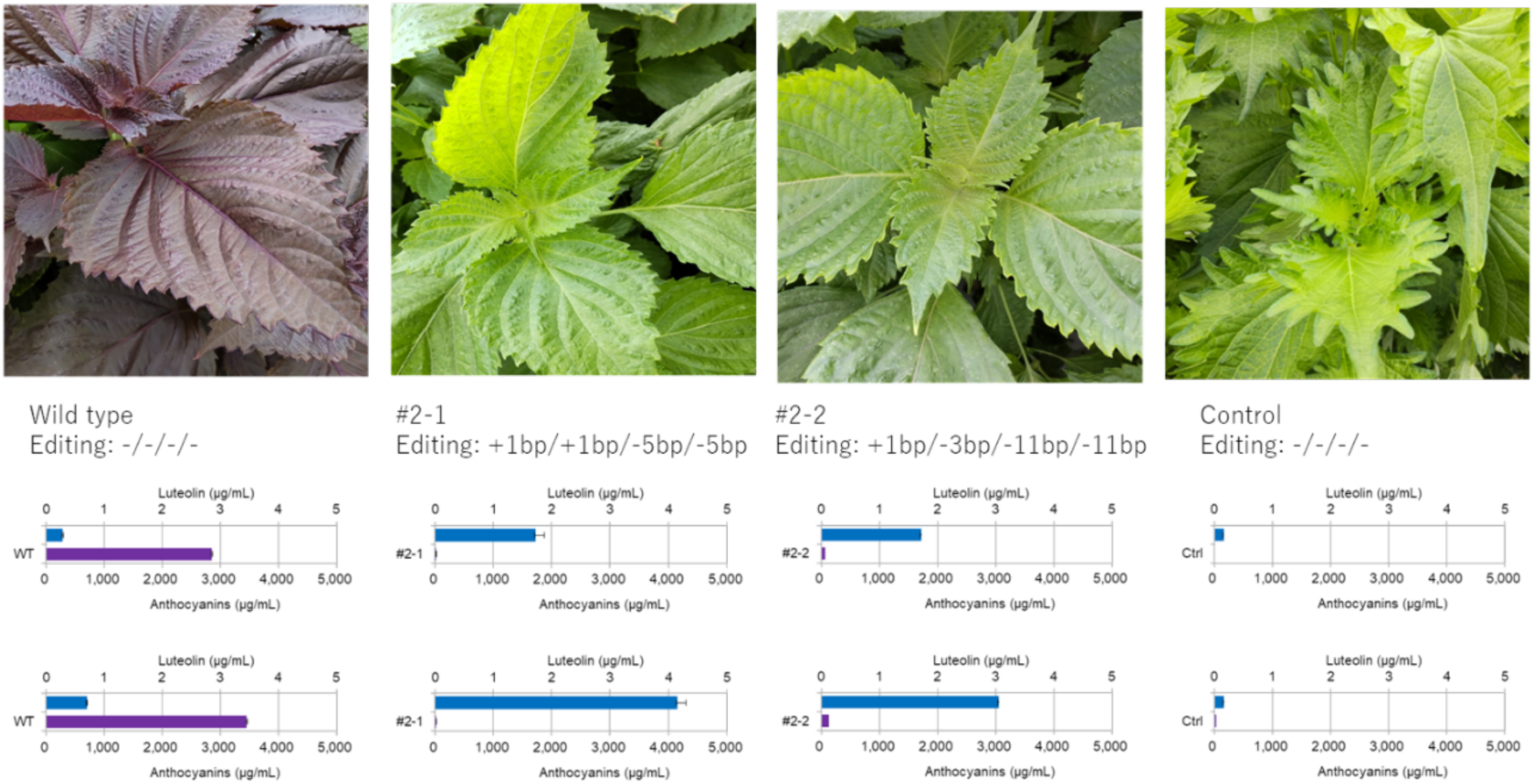
Anthocyanin and luteolin contents in the T₁ generation. Sampling was conducted on June 26 (upper panel) and September 11 (lower panel). Data are expressed in terms of fresh weight (per g FW).

In WT plants, luteolin levels were only 0.3 µg/g FW in June and 0.7 µg/g FW in September, whereas the five genome-edited lines accumulated 1.0–2.2 µg/g FW and 3.0–4.2 µg/g FW, respectively. For example, line #2-1 accumulated 1.7 µg/g FW in June and 4.2 µg/g FW in September, representing an approximately six-fold increase relative to WT (Fig. 2). This trend was consistently observed across all five T₁ individuals, with significant increases (P < 0.001; except for #2-3 in June, P < 0.05) (Fig. S2).

Rosmarinic acid content was consistently elevated in edited lines relative to WT in both June and September, particularly in #2-1 (up to 10,125 µg/g FW in September), suggesting redirection of metabolic flux toward other phenolic compounds. Perillaldehyde levels were more variable, with some lines showing higher concentrations than WT, and others showing lower concentrations, indicating that this monoterpenoid is less directly affected by *F3H* editing and may also be influenced by developmental stage or environmental conditions.

### Transcriptomic analysis of flavonoid pathway genes in *F3H*-edited lines

Transcriptome analysis revealed that *F3H* expression was markedly reduced, whereas the expression of *PAL*, *C4H*, *CHI*, and several *F3’H* family genes was upregulated (Fig. 3; Table S2), supporting a metabolic shift that redirects naringenin flux toward luteolin biosynthesis.

**Fig. 3.**
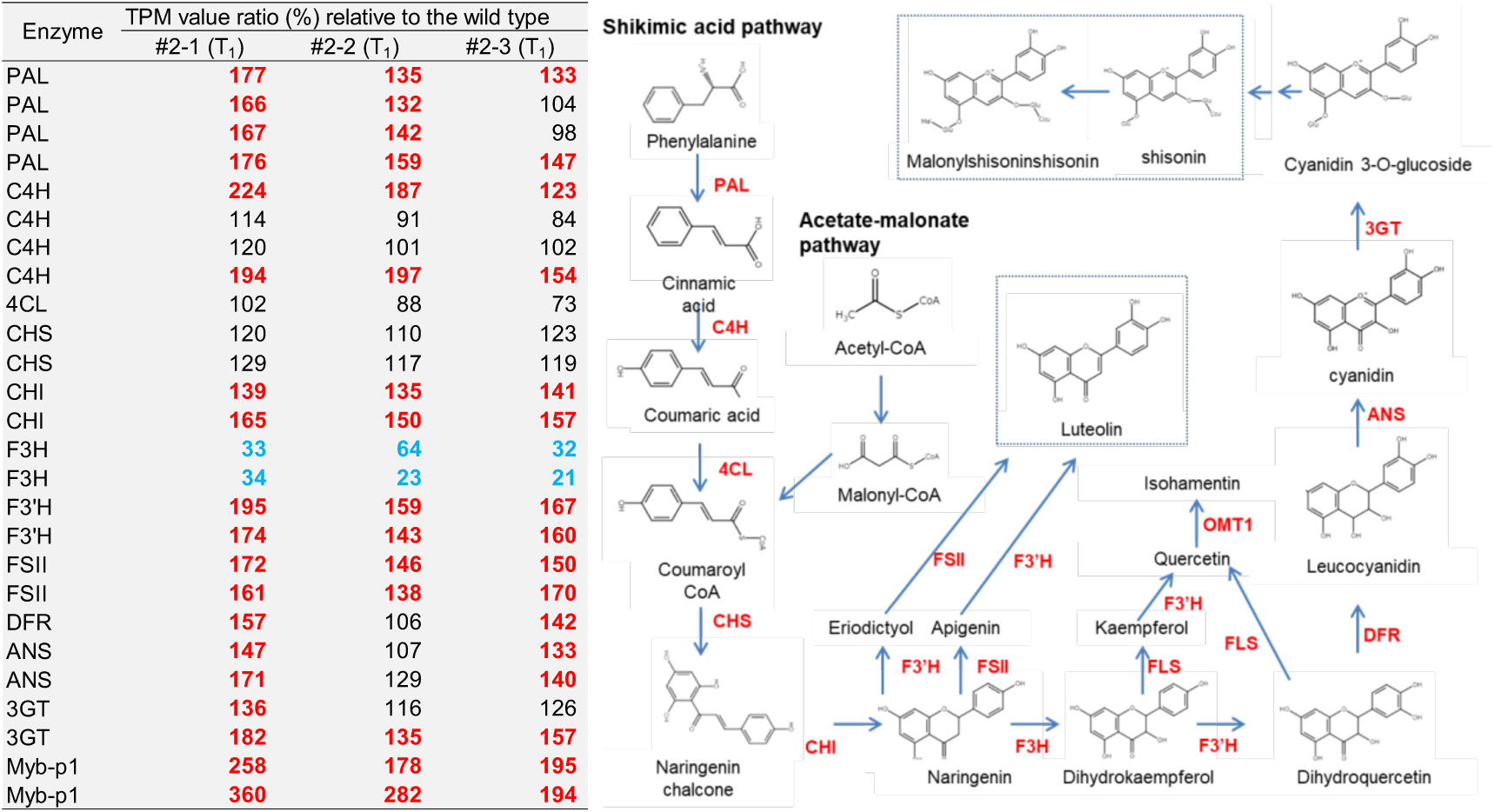
Major anthocyanin biosynthetic pathway and expression profile of key biosynthetic genes in P. frutescens. Genes listed in Table S2 with TPM values of 50 or higher are shown. Numerical values indicate the relative expression levels (%) compared to the wild type. Genes with expression levels increased by more than 30% are highlighted in bold red, while those with decreased expression are shown in bold blue. PAL, phenylalanine ammonialyase; C4H, cinnamic acid 4-hydroxylase; 4CL, 4-coumaric acid: CoA ligase; ACC, acetyl-CoA carboxylase; CHS, chalcone synthase; CHI, chalcone isomerase; F3H, flavanone 3-hydroxylase; F3’H, flavonoid 3’-hydroxylase; FSII, flavonol synthase II; FLS, flavonol synthase; OMT1, O-methyltransferase 1; DFR, dihydroflavonol 4-reductase; ANS, anthocyanidin synthase; 3GT, UDP-glucose: anthocyanidin 3-O-glucosyltransferase.

**Fig. 4.**
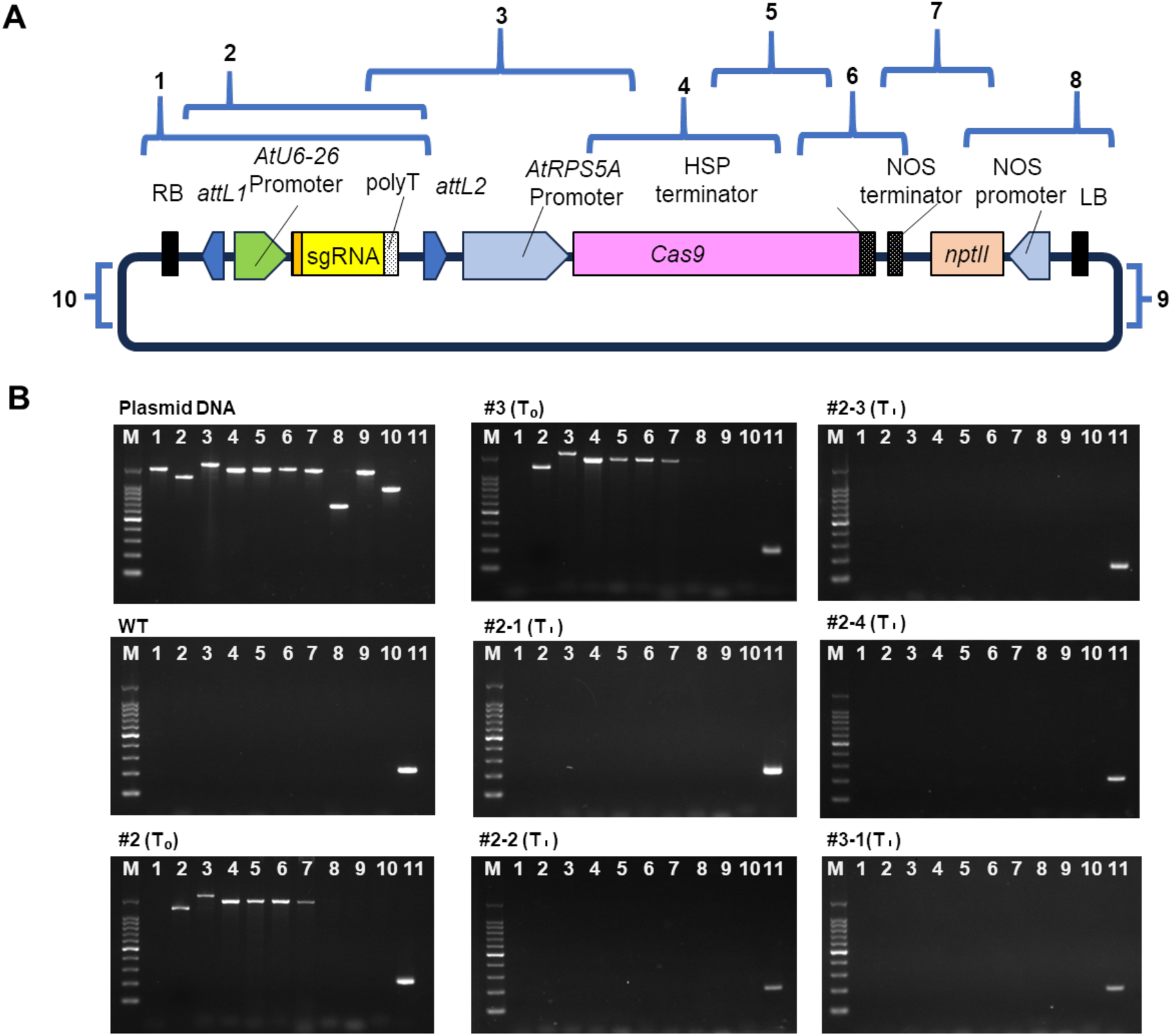
PCR analysis of putative null segregants. Primer sequences and expected amplicon sizes are listed in Table S1. **(A)** Schematic diagram of the HEMCas9-nptII_AtU6gRNA2 construct. The plasmid is divided into regions 1–10, of which regions 2–7 are located within the T-DNA region. **(B)** PCR amplification results. Lane M: 100 bp DNA ladder; the top band corresponds to 1500 bp. Lanes 1–10: PCR amplification of genomic/plasmid DNA corresponding to regions 1–10 in panel (A), used to detect presence or absence of plasmid sequences. Lane 11: PCR amplification of the endogenous *F3H* gene as a positive control for plant genomic DNA.

Upstream enzymes of the phenylpropanoid pathway, including *PAL* and *C4H*, tended to be expressed at higher levels in the edited lines compared with their expressions in the WT, suggesting that metabolic flux may have been redirected upstream because of constraints in downstream flavonoid biosynthesis.

Among flavonoid biosynthetic genes, *CHS* and *CHI* maintained expression levels at 110–165% of WT across all edited lines. In contrast, *F3H* transcripts were substantially reduced, reaching only 21–64% of WT, confirming effective loss of *F3H* activity. Notably, *F3’H* expression increased to 143–195% of WT, consistent with the observed elevation of luteolin and luteolin-related compounds.

Downstream flavonoid enzymes, including *DFR* and *ANS*, also exhibited expression levels of 106–171% of WT, indicating that partial pathway activity was retained despite the metabolic bottleneck caused by *F3H* disruption.

O-methyltransferase (*OMT3*) showed moderate upregulation (approximately 100–130% of WT), although its absolute transcript abundance was low, potentially contributing to the increased accumulation of methylated flavonoid derivatives.

Furthermore, transcription factors involved in flavonoid biosynthesis, particularly *Myb-p1*, were strongly upregulated in T₁ individuals (178–360% of WT), suggesting activation of regulatory networks in response to *F3H* impairment.

### Generation of genome editing null-segregant plants

To obtain genetically stable genome-edited plants, it was necessary to remove all genome editing components from the host genome. Segregation of vector-positive and -negative progeny within each T₁ family followed the expected 3:1 Mendelian ratio, as supported by chi-square (χ²) tests (p > 0.05) (Table S3). These results indicate that the T-DNA was inserted at a single genomic locus in each line.

PCR assays were further performed on T₁ individuals (#2-1, #2-2, #2-3, #2-4, and #3-1) using primer sets spanning the entire vector backbone (Table S4). While all target regions were successfully amplified using plasmid HEMCas9-nptII_AtU6gRNA2 as a positive control, no amplification was detected in any T₁ individual except at the endogenous F3H locus (Fig. 3), thus confirming the absence of vector-derived sequences in these plants.

To additionally evaluate potential residual vector fragments, RNA sequencing (RNA-seq) data were screened using K-mer analysis (k = 31). A short sequence identical to a vector-derived fragment was detected at a low level in both WT and line #2-1 (WT: GTTAAAATAAGGCTAGTCCGTTATCAACTTG; #2-1: AGCAAGTTAAAATAAGGCTAGTCCGTTATCA). However, these sequences were also detected in the WT samples and did not map to the reference genome (Pfru_yukari_1.0). BLASTn analysis revealed high similarity to *Escherichia coli* strain K12 (AP027461.1), suggesting that these reads likely originated from minor experimental contamination rather than from plant genomic DNA.

Taken together, these results provide strong evidence that the T₁ individuals (#2-1, #2-2, #2-3, #2-4, #3-1) are null segregants completely free of genome editing components.

## Discussion

In the present study, we successfully generated green *Perilla* lines by genome editing of the *F3H* gene, a key enzyme in the anthocyanin biosynthetic pathway of red perilla. The edited plants exhibited substantially reduced anthocyanin accumulation and increased luteolin levels. These findings demonstrate the central role of *F3H* in anthocyanin production in perilla and indicate that its disruption redirects metabolic flux toward the bioactive flavonoid luteolin. The resulting metabolic and transcriptional changes provide the first direct physiological evidence for *F3H* function in perilla. More broadly, the present study illustrates the utility of genome editing for functional genomics in non-model species and highlights its potential for developing high-value crops enriched in health-promoting metabolites.

Anthocyanins are major pigments distinguishing red and green perilla, and their biosynthetic pathway has been well characterized (Saito & Yamazaki, 2002; Fukushima et al., 2015). Recent genetic studies identified *F3H*, *C4H1*, and the transcription factor *MYB113b* as major determinants of anthocyanin production in perilla (Xie et al., 2025). Consistent with observations in *Arabidopsis thaliana*, where *F3H* loss-of-function mutants alter flavonoid accumulation (Pelletier et al., 1999), disruption of *F3H* in perilla is expected to block the conversion of naringenin to dihydrokaempferol and consequently reduce anthocyanin biosynthesis. This metabolic bottleneck likely enhances flux toward luteolin biosynthesis through intermediates such as eriodictyol and apigenin, mediated by *F3’H* and *FSII* activity (Fig. 3). Overall, the results of the present study indicate that disruption of *F3H* not only suppressed anthocyanin biosynthesis but also resulted in a reorganization of the transcriptional landscape of the phenylpropanoid and flavonoid pathways, enhancing metabolic flux toward luteolin and other flavonoid derivatives.

Metabolite profiling revealed a distinct metabolic shift characterized by reduced anthocyanin levels and increased luteolin accumulation (Fig. 2, Fig. S2). HPLC analysis detected luteolin and its analogs as a single unresolved peak that could not be chromatographically resolved under the conditions used, suggesting possible co-elution of structurally related compounds. HPLC–TOF–MS analysis further identified an unknown metabolite with a molecular weight of 300.0634 (C₁₆H₁₂O₆), consistent with a mono-methylated derivative of luteolin (molecular weight 286.0477, C₁₅H₁₀O₆). O-methylation is known to increase flavonoid lipophilicity and may enhance metabolite stability during intracellular transport or storage (Walle, 2009; Koirala et al., 2016). Several methylated luteolin derivatives, including chrysoeriol and diosmetin, which are 3′- and 4′-O-methylated derivatives of luteolin, respectively, have been reported to accumulate in perilla seeds (Lee et al., 2013; Lee et al., 2017a; Kim & Lee, 2019). However, the chromatographic retention time of the unknown compound detected in this study did not match that of authentic diosmetin. The MS/MS fragment ion pattern derived from the precursor ion at m/z 299.0556 did not match that of chrysoeriol (Lee et al., 2013; Lee et al., 2017b), but showed high similarity to that of hispidulin, although the chromatographic retention time differed from that of the authentic standard. These observations suggest that the detected metabolite is not diosmetin, chrysoeriol or hispidulin itself, but a distinct methylated luteolin-derived flavone isomer with a similar core structure.

O-methylation of flavonoids is catalyzed by O-methyltransferases *(OMT*s) using S-adenosylmethionine as the methyl donor. In perilla, *PfOMT3* has been cloned and shown to catalyze O-methylation at the 7-OH position of certain flavonoid substrates (Park et al., 2020). Although luteolin has been reported as a substrate of *PfOMT3*, the precise site of O-methylation has not been experimentally determined. The positional specificity exhibited by *PfOMT3* toward other flavonoid substrates may provide useful clues to its potential role in flavonoid O-methylation. In tomato, the *OMT3* homolog *SlOMT3* has been reported to convert luteolin to chrysoeriol (Cho et al., 2012). Indeed, multiple *PfOMT3* homologs are present in the perilla genome (Table S2), and functional diversification among these enzymes may contribute to the production of structurally distinct methylated flavones. On the other hand, the Arabidopsis enzyme *AtOMT1* has also been reported to catalyze O-methylation of luteolin (Muzac et al., 2000; Tohge et al., 2007), indicating that enzymes other than *PfOMT3* and its homologs may be involved in luteolin O-methylation.

To definitively determine the methylation position and the enzymatic basis of this compound, further integrative metabolomic and genomic analyses will be required. Although LC–MS/MS analysis provided strong evidence that the detected compound represents a methylated luteolin-derived flavone isomer, definitive determination of the methylation position will require structural elucidation by NMR spectroscopy, which was beyond the scope of the present study. Moreover, the involvement of O-methyltransferases other than *PfOMT3* and its homologs cannot be excluded, and further studies will be needed to clarify this point. In addition, the present study was limited to metabolite profiling at a specific developmental stage, and further temporal and tissue-specific analyses will be required to fully elucidate the regulatory network.

RNA-seq analysis revealed strong downregulation of *F3H* transcripts, likely resulting from nonsense-mediated mRNA decay triggered by frameshift mutations in exon 1 (Drechsel et al., 2013; Tuladhar et al., 2019; Luha et al., 2024). Notably, even lines lacking functional *F3H* accumulated trace anthocyanins, suggesting the presence of residual *F3H* activity at another genomic locus or the involvement of an alternative enzyme with weak C3-hydroxylation activity. Other anthocyanin biosynthetic genes—including *CHI*, *PAL*, *C4H*, *F3’H*, *FSII*, and *3GT*—were upregulated in the edited lines. These genes are normally coordinately activated in red perilla (Saito and Yamazaki, 2002; Fukushima et al., 2015). Their elevated expression likely reflects the absence of anthocyanin accumulation, which may relieve feedback inhibition and maintain pathway activation. The transcription factor *Myb-p1*, which regulates *DFR* expression in a light-dependent manner (Gong et al., 1999b), was also strongly upregulated. By contrast, *MYB113b*, which activates *C4H* and *F3H* (Xie et al., 2025), remained weakly expressed. Similar to *A. thaliana*, where multiple MYB groups regulate anthocyanin biosynthesis (Stracke et al., 2007; Dubos et al., 2010), perilla may employ different MYB modules depending on genotype or tissue type, warranting further investigation. Indeed, recent studies in diverse plant species have highlighted functional diversification of MYB transcription factors in the regulation of anthocyanin and flavonoid biosynthesis (Li et al., 2022; Menconi et al., 2023). These observations warrant further investigation into the roles of MYB regulators in *P. frutescens*.

Disruption of *F3H* likely caused a redistribution of metabolic flux within the phenylpropanoid pathway. Notably, rosmarinic acid levels were significantly elevated in the edited lines (Fig. S2). Flavonoids and other phenolic compounds, including rosmarinic acid, share the common precursor p-coumaroyl-CoA (Jakovljević et al., 2025; Petersen et al., 2009). Upregulation of upstream genes such as PAL and C4H may have increased the supply of this precursor, thereby contributing to the enhanced accumulation of rosmarinic acid (Vogt, 2010). These findings suggest that targeted modification of a single flavonoid biosynthetic enzyme can substantially influence the broader phenolic metabolite network in perilla. However, because enzyme activities and metabolic fluxes were not directly measured in the present study, the causal relationship between the upregulation of PAL and C4H and the enhanced accumulation of rosmarinic acid requires further verification through biochemical analyses.

Genome editing enables the development of cultivars that were previously unattainable. Because F3H exhibits incomplete dominance, crosses between red and green perilla yield lines with intermediate anthocyanin levels (Kinoshita et al., 2023; Xie et al., 2025), but increasing luteolin accumulation is not straightforward. The −3 bp deletion allele generated in the present study retains partial enzyme activity and thus represents an intermediate allele that allows stepwise control of pigmentation and luteolin content, serving as a valuable new genetic resource. In the edited lines generated in the present study, PCR and k-mer analysis of RNA-seq data confirmed the absence of vector-derived sequences in T₁ individuals, suggesting a low likelihood of classification as genetically modified under regulatory frameworks such as the Cartagena Protocol. Nevertheless, for future commercialization, evaluation in accordance with the regulations of individual countries will be required.

Overall, the present study demonstrates that targeted disruption of *F3H* modulates the flavonoid pathway, generating *Perilla* lines with strongly reduced anthocyanin levels and enhanced luteolin content. This work provides a molecular framework for tailoring the metabolic composition of perilla and highlights genome editing as a promising strategy for developing functional crops with improved health benefits and commercial value.

## Experimental Procedures

### Plant materials and growth conditions

For genome editing, *P. frutescens* var. *crispa* cv. ‘Hoko-3’ was used. The transgenic T₀ plants generated as described below were acclimatized and subsequently grown in a plant growth chamber. The plants were initially cultivated under long-day (LD) conditions (photosynthetic photon flux density, PPFD, 100 μmol m⁻² s⁻¹; 16 h light/8 h dark) at 25 °C for approximately 10 weeks, after which they were shifted to short-day (SD) conditions (PPFD 100 μmol m⁻² s⁻¹; 8 h light/16 h dark) to induce flowering and self-pollination for seed production. A portion of the harvested seeds was grown under LD conditions in a growth chamber as the T₁ generation. Among them, null segregants identified as described below were propagated by excising axillary shoots and rooting them in rockwool to obtain multiple clonal plants. These genome-edited lines, together with the commercial green perilla cultivar ‘Ooba Ao-shiso’ (Tohokuseed Co., Utsunomiya, Ltd., Japan) as a control, were transferred to an isolated greenhouse and cultivated hydroponically under natural temperature conditions (20–40 °C) from May to October.

### Vector construction

At the time of this study, the genome sequence of ‘Hoko-3’ had not been released. Therefore, the *F3H* gene sequence was assembled, and a gRNA was designed within the first exon (Supplementary Material 1, Fig. S1). Oligonucleotides synthesized with overhangs compatible with the gRNA and restriction enzyme sites were annealed to form double-stranded DNA and cloned into the entry vector pENTR_ATU6 (Nobusawa et al., 2021) via *BbsI* restriction sites. The gRNA expression cassette was subsequently transferred into HEMCas9-nptII (formerly pGWB401_AtRPS5A-Cas9) (Yamatani et al., 2022) by an LR Clonase reaction (Fig. 1A). This binary vector was derived from pGWB401, which was developed based on the Gateway binary vector system (Nakagawa et al., 2007). In this construct, Cas9 is driven by the RPS5A promoter and terminated by the HSP terminator. The resulting plasmid was introduced into *Agrobacterium tumefaciens* strain EHA105 by electroporation.

### Agrobacterium*-*mediated transformation

Seeds of *P. frutescens* cv. ‘Hoko-3’ were surface-sterilized by immersion in 70% ethanol for 1 min, followed by treatment with 1% sodium hypochlorite solution for 15 min, and rinsed four times with sterile distilled water. Sterilized seeds were sown on half-strength Murashige and Skoog (MS) medium (Murashige and Skoog, 1962) supplemented with 3% sucrose and solidified with 0.3% gellan gum. After germination, seedlings were grown under long-day (16 h light/8 h dark) conditions at 23 °C for 1 week, and hypocotyls (n = 239) were excised to a length of 0.5–0.8 cm. *Agrobacterium tumefaciens* strain EHA105 was cultured in YEP medium (10 g L⁻¹ yeast extract, 10 g L⁻¹ peptone, and 10 g L⁻¹ NaCl) at 28°C with shaking at 100 rpm for 24 h. Bacterial cells were collected by centrifugation at 2,000 × g for 10 min using a refrigerated centrifuge(LCX-100, TOMY Seiko Co., Ltd., Tokyo, Japan), and resuspended in an infiltration buffer containing 0.1 mmol L⁻¹ acetosyringone, 10 mmol L⁻¹ MgCl₂, and 1% Tween 20, and the optical density at 600 nm (OD₆₀₀) was adjusted to 0.1. Hypocotyl segments were immersed in the suspension for 30 min to facilitate infection. Following infection, explants were co-cultivated for 3 days at 23 °C on half-strength MS solid medium supplemented with 1.0 mg L⁻¹ 6-benzylaminopurine (BA) and 0.1 mg L⁻¹ α-naphthaleneacetic acid (NAA). They were then transferred to a decontamination medium containing 500 mg L⁻¹ cefotaxime and incubated under the same conditions for 7 days. Subsequently, explants were transferred to a selection medium containing 50 mg L⁻¹ kanamycin, with subculture being carried out every 2 weeks for callus induction and selection. After 6–10 weeks, shoots regenerated from the calli were excised and transferred to hormone-free half-strength MS solid medium for root induction and plant regeneration.

### Confirmation of transformation

PCR analysis confirmed the presence of T-DNA-derived sequences in regenerated plants. Genomic DNA was extracted from leaves using the ISOSPIN Plant DNA Kit (Nippon Genetics, Tokyo, Japan). PCR amplification was performed with primers designed based on the T-DNA sequence (Table S3) and Quick Taq® HS DyeMix (Toyobo, Osaka, Japan). The PCR conditions were as follows: an initial denaturation step at 95 °C for 5 min, followed by 30 cycles at 95 °C for 30 s, 55 °C for 30 s, and 72 °C for 30 s, with a final extension at 72 °C for 30 s.

### Genome editing validation

Genome editing events were verified by amplicon sequencing. Primers containing linker, tag, and target sequences were used to amplify the target region simultaneously (Table S3). Genomic DNA extracted from leaves of T₁ plants was used as template, and PCR was performed under the following conditions: initial denaturation at 95 °C for 5 min, followed by 30 cycles of 95 °C for 30 s, 55 °C for 30 s, and 72 °C for 30 s, with a final extension at 72 °C for 30 s. PCR products were confirmed by electrophoresis, and their concentrations were estimated. All PCR products were pooled, purified with Ampure XP (Beckman Coulter, Brea, CA, USA), and subjected to paired-end sequencing (300 bp) on an Illumina MiSeq platform by Hokkaido System Science Co., Ltd. (Hokkaido, Japan).

### Identification of null segregants

For T₁ plants, PCR screening was performed as described for T₀ plants. In five individuals cultivated in an isolated greenhouse (#2-1, #2-2, #2-3, #2-4, and #3-1), the persistence of vector sequences was examined using primers covering the entire vector backbone (Table S1; Fig. 2) (Nishihara et al., 2023). In addition, the transcriptome RNA-seq data, which were also used for subsequent gene expression analyses, were employed here to examine the presence of vector-derived sequences using a K-mer–based approach. K-mers of length 31 bp were first generated from the vector sequence using Jellyfish (ver. 2.3.0), and perfectly matching reads were extracted from each RNA-seq dataset. K-mers were subsequently re-extracted from the matched reads, and sequences shared with the vector were identified. These sequences were mapped to the reference genome of ‘Hoko-3’ (Pfru_yukari_1.0; https://perilla.annotation.jp/) using Bowtie2 (ver. 2.4.2) (Langmead & Salzberg, 2012). Finally, BLASTn searches against the NCBI nonredundant nucleotide database were conducted to ensure that no nonspecific sequences remained (Camacho et al., 2009).

### Metabolite profiling by HPLC and LC-QTOF/MS

Upper leaves of hydroponically grown plants in an isolated greenhouse were collected from multiple clonal individuals on June 26 and September 11. For the analysis of perillaldehyde and rosmarinic acid, 0.5 g of fresh leaves was chopped and extracted overnight at 10 °C in 5 mL of 80% methanol. The extract was then subjected to sonication for 10 min and centrifugation at 670 × g for 10 min using a refrigerated centrifuge (Model 5420, Kubota Corporation, Tokyo, Japan), the supernatant was subsequently collected. Anthocyanins were extracted using the same procedure except that 0.5% trifluoroacetic acid (TFA) was used as the extraction solvent. For the analysis of luteolin and its related compounds, 1 g of leaves temporarily frozen at –20 °C was chopped, extracted with 5 mL of 80% methanol, sonicated for 15 min, and centrifuged under the same conditions; the resulting supernatant was collected. All analyses were performed with three biological replicates. Quantification of perillaldehyde, rosmarinic acid, anthocyanins, luteolin, and its derivatives was conducted on a Shimadzu HPLC system (LC-20AT pump, SPD-20AV UV detector, CBM-20A lite system controller; Shimadzu Corporation, Kyoto, Japan) under the conditions listed in Table S5. Detailed analysis of luteolin was conducted using a liquid chromatography–quadrupole time-of-flight mass spectrometry (LC-QTOF-MS) system, comprising a 1260 Infinity II HPLC system (Agilent Technologies, Santa Clara, CA,USA) coupled with an X500R QTOF mass spectrometer (AB Sciex LLC, Framingham, MA, USA), under the conditions described in Table S6.

### Transcriptome analysis

Transcriptome analysis was performed to evaluate gene expression levels in genome-edited plants and to confirm the absence of vector-derived sequences within expressed genes. Total RNA was extracted from leaves harvested on June 18 from hydroponically grown wild-type and three T₁ plants (#2-1, #2-2, and #2-3) cultivated in an isolated greenhouse using the ISOSPIN Plant RNA Kit (NIPPON GENE Co., Ltd., Tokyo, Japan). RNA-seq libraries were constructed with the VAHTS Universal V8 RNA-seq Library Prep Kit for Illumina. Paired-end sequencing (2 × 150 bp) was carried out on a NovaSeq 6000 platform (Illumina, San Diego, CA, USA) at a depth of 6 GB per sample by Azenta Japan (Tokyo, Japan). Raw reads were processed using Trimmomatic (ver. 0.39) to remove adapter sequences and low-quality bases. Clean reads were mapped to the ‘Hoko-3’ reference genome using HISAT2 (ver. 2.2.1) (Kim et al., 2019b), and transcript abundance was quantified with StringTie (ver. 2.1.7) (Pertea et al., 2015).

### Accession number

The RNA-seq data generated in this study have been deposited in the DDBJ BioProject database under the accession number PRJDB38062

## Acknowledgements

The plasmids pENTR_ATU6 and HEMCas9-nptII were generously provided by the National Bioresource project Chrysanthemums. We would like to thank Editage (www.editage.jp) for English language editing.

## List of Abbreviations

T₀: Transgenic generation 0 (plants regenerated immediately after transformation)
T₁: Transgenic generation 1(Progeny of T₀ plants obtained by self-pollination)
PAL: phenylalanine ammonialyase
C4H: cinnamic acid 4-hydroxylase
4CL: 4-coumaric acid: CoA ligase
CHS: chalcone synthase
CHI: chalcone isomerase
F3’H: flavonoid 3’-hydroxylase
FSII: flavone synthase I
F3H: flavanone 3-hydroxylase
FLS: flavonol synthase
OMT1: O-methyltransferase 1
DFR: dihydroflavonol 4-reductase
ANS: anthocyanidin synthase
3GT: UDP-glucose: anthocyanidin 3-O-glucosyltransferase
WT: Wild type
RNA: sequencing (RNA-seq)
LD: Long day
SD: Short day
PPFD: Photosynthetic photon flux density

## Short legends for Supporting Information

Supplementary Methods S1. Target gene sequence identification and gRNA design for CRISPR–Cas9 genome editing in *P. frutescens*.

Supplementary Figure S1. Sequence organization of the *P. frutescens F3H* gene for gRNA design.

Supplementary Figure S2. Metabolite accumulation in *P. frutescens* collected on June 26 and September 11.

Supplementary Table S1. Genome editing outcomes for the *F3H* gene.

Supplementary Table S2. List of enzyme-coding genes involved in the representative anthocyanin biosynthetic pathway of *P. frutescens*.

Supplementary Table S3. Segregation of transgene in T₁ progeny. Supplementary Table S4. List of Oligonucleotides used in this study.

Supplementary Table S5. HPLC conditions for the quantification of major compounds. Supplementary Table S6. Analytical conditions for luteolin quantification using LC-QTOF/MS.

## Author Contributions

SM performed RNA and DNA extractions, PCR analyses, microbial transformation, indoor cultivation, transcriptome analysis, and wrote the manuscript. MN constructed the plasmids and revised the manuscript. SC and MK conducted cultivation, plant transformation, and growth in a controlled environment facility. AF carried out RNA and DNA extractions and PCR analyses. CN and TI were responsible for sampling and component analysis using HPLC. JK performed component analysis using LC-QTOF-MS. KT conducted transcriptome analysis, critically reviewed the manuscript, and contributed to manuscript revision. HB conceived the research plan, critically reviewed the manuscript, and provided final approval.

## Funding

This work was supported by the Hiroshima Prefectural Government “Hiroshima Health and Medical-Related Industry Creation Support Project” and the Center of Innovation for Bio-Digital Transformation (BioDX), an open innovation platform for industry-academia co-creation of JST (COI-NEXT, JPMJPF2010).

## Conflict of interest statement

The authors declare no conflict of interest.

## Notes

### Competing Interest Statement

The authors have declared no competing interest.

