## Supporting information for "Genome editing of the flavanone 3-hydroxylase gene produces a green *Perilla* line with elevated luteolin accumulation"

**Title**

**Content**

Supplementary Method: S1

Supplementary Figures: S1−S2

Supplementary Tables: S1−S6

### Supplementary Method

**Method S1. Target gene sequence identification and gRNA design for CRISPR–Cas9 genome editing in *P. frutescens*.**

To perform genome editing using CRISPR–Cas9, it is necessary to design gRNAs specific to the target gene. At the start of this study, the genome sequence of Perilla frutescens var. japonica 'Hoko-3' had not yet been determined; therefore, RNA-seq and Sanger sequencing were conducted to obtain sequence information.

Total RNA was extracted by Bioengineering Laboratory Co., Ltd. (Kanagawa, Japan), and approximately 3 Gb of 151 bp paired-end sequencing data were generated using the Illumina NextSeq 500 platform. The obtained reads were processed with Trimmomatic (v0.39) to remove low-quality reads and adapter sequences (Bolger et al., 2014), followed by de novo assembly using Trinity (v2.14.0) (Grabherr et al., 2011). Coding regions were predicted using TransDecoder (v5.5.0).

The F3H CDS (Accession No. AB000286.1) was used as a query for local BLAST searches, and representative sequences showing homology were extracted. The extracted sequences were aligned using ClustalW, as shown in Fig. S1A.

Based on these sequences, primers for sequencing were designed, and Sanger sequencing was performed (Table S1, Fig. S1). From the resulting genomic sequences, the first exon was predicted, and allele information in the allotetraploid genome was inferred from the sequencing traces. Considering that single-nucleotide mismatches located more than 13 bp away from the PAM do not significantly affect genome editing efficiency (Fu et al., 2013; Hsu et al., 2013; Anderson et al., 2015), gRNAs were designed accordingly (Fig. S1). Although multiple gRNA candidates were designed, only Target1 was used in subsequent experiments.

**Supplementary Figures**

**A**

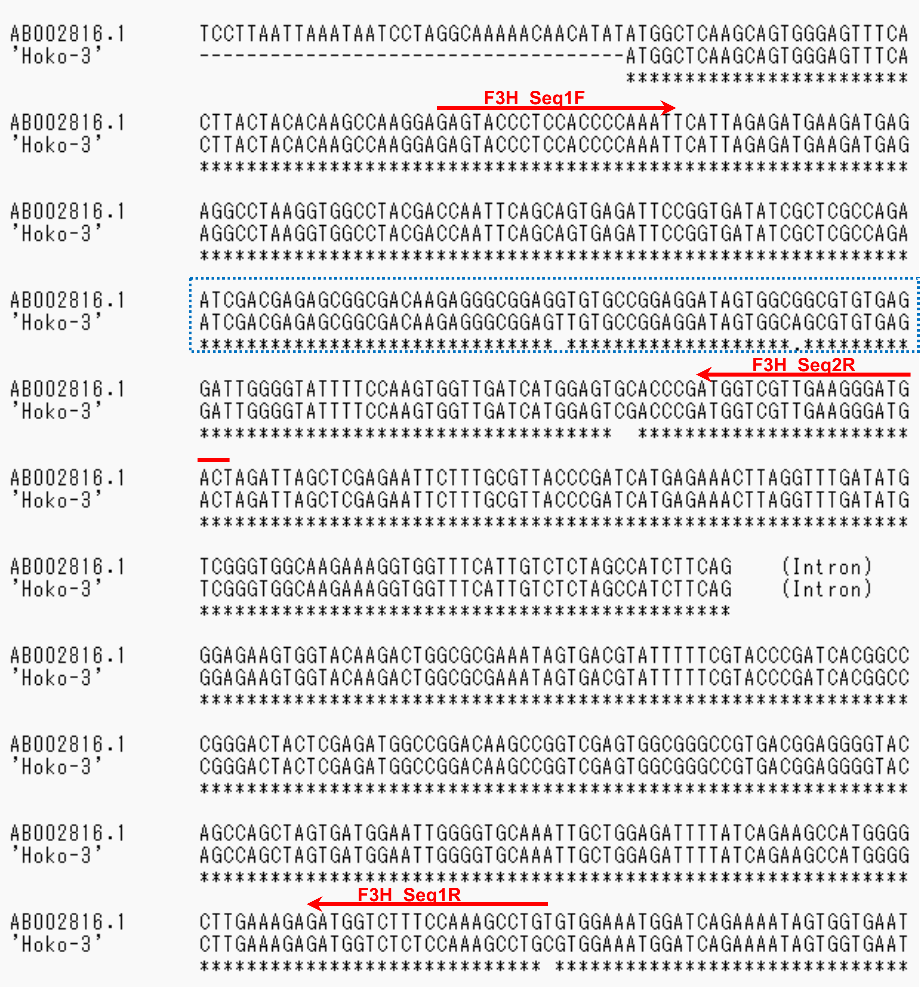


**B**

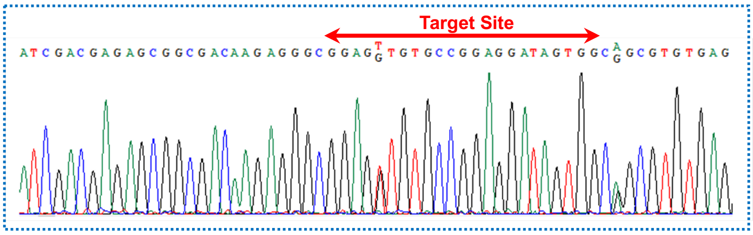


**Fig. S1. Sequence organization of the *P. frutescens F3H* gene for gRNA design.**

(A) Alignment of the CDS of the perilla *F3H* gene with the 'Hoko-3' sequence reconstructed from RNA-seq data. Red arrows indicate the primers used for sequencing.

(B) Sanger sequencing of 'Hoko-3' in the region highlighted by the blue dashed box in panel (A). Red arrows indicate the target site and PAM sequence.

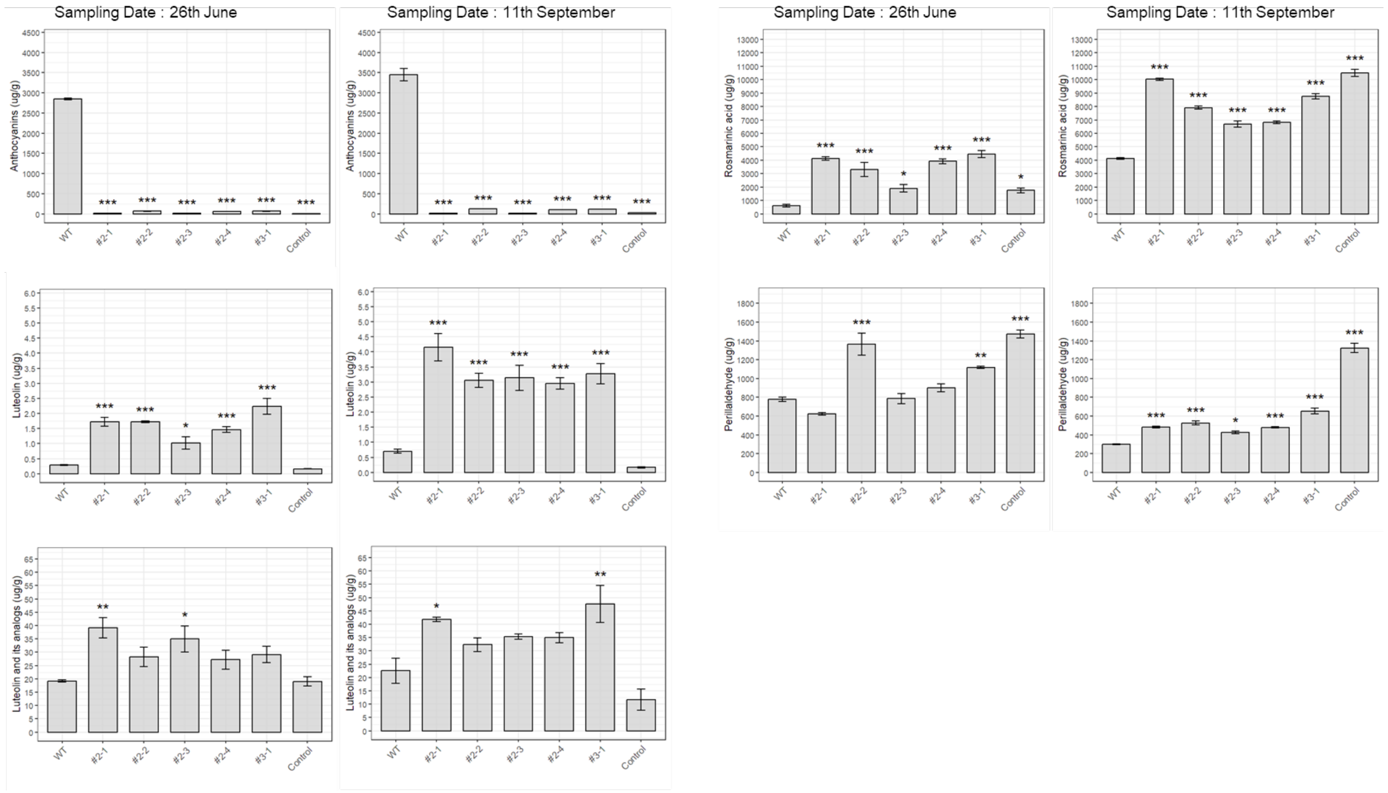


**Fig. S2. Metabolite accumulation in *P. frutescens* collected on June 26 and September 11.**
Data from June 26 and September 11 are shown in the left and right panels, respectively. Quantification of five major metabolites—Anthocyanins, Luteolin, Luteolin and its analogs, Rosmarinic acid, and Perillaldehyde—in wild-type *P. frutescens* 'Hoko-3' (WT, red perilla), genome-edited lines (#2-1, #2-2, #2-3, #2-4, #3-1), and the non-red cultivar 'Ooba Ao-shiso' (Control, green perilla). Bars indicate the mean ± SE of three biological replicates (n = 3). Statistical analyses were performed separately for each compound and sampling date using one-way ANOVA followed by Dunnett's test, with WT ('Hoko-3') used as the reference group. Asterisks denote significant differences relative to WT (*P < 0.05; **P < 0.01; ***P < 0.001).

### Supplementary Table

**Table S1. Genome editing outcomes for the *F3H* gene.**

| Line (Genelation) | Allele (Chromosome) | Target Region | In/del |
|---|---|---|---|
| Wild type | 1 (Pfg07) | GGCGGAGGTGTGCCGGAGGATAGTGGCGGCGTGTGA | - |
| | 1 (Pfg19) | GGCGGAGTTGTGCCGGAGGATAGTGGCAGCGTGTGA | - |
| #1 ($T_0$) | 1 (Pfg07) | GGCGGAGGTGTGCC------TAGTGGCGGCGTGTGA | -6bp |
|  | 2 (Pfg07) | GGCGGAGG-----------------------TGTGA | -23bp |
| | 3 (Pfg07) | GGCGGAGGTGTGCCGGAGGATTAGTGGCGGCGTGTGA | +1bp |
|  | 4 (Pfg07) | GGC--------------------TGGCGGCGTGTGA | -20bp |
| | 1 (Pfg19) | GGCGGAGTTGTGCCGGAGGAATAGTGGCAGCGTGTGA | +1bp |
| | 2 (Pfg19) | GGCGGAGTTGTGCCGGAGG-TAGTGGCAGCGTGTGA | -1bp |
| #2 ($T_0$) | 1 (Pfg07) | GGCGGAGGTGTGCCGGAGG-TAGTGGCGGCGTGTGA | -1bp |
| | 2 (Pfg07) | GGCGGAGGTGTGCCG---GATAGTGGCGGCGTGTGA | -3bp |
|  | 1 (Pfg19) | GGCGGAGTTGTGCCG-----TAGTGGCAGCGTGTGA | -5bp |
|  | 2 (Pfg19) | GGCGGAGTT-----------TAGTGGCAGCGTGTGA | -11bp |
| #3 ($T_0$) | 1 (Pfg07) | GGCGGAGGTGTGCCGGAGGA--GTGGCGGCGTGTGA | -2bp |
| | 2 (Pfg07) | GGCGGAGGTGTGCCG---GATAGTGGCGGCGTGTGA | -3bp |
|  | 1 (Pfg19) | GGCGGAGTTGTGCCG-----TAGTGGCAGCGTGTGA | -5bp |
|  | 2 (Pfg19) | GGCGGAGTT-----------TAGTGGCAGCGTGTGA | -11bp |
| #4 ($T_0$) | 1 (Pfg07) | GGCGGAGGTGTGCCGGAGGAATAGTGGCGGCGTGTGA | +1bp |
| | 1 (Pfg19) | GGCGGAGTTGTGCCGGAGGAATAGTGGCAGCGTGTGA | +1bp |
| #2-1 ($T_1$) | 1 (Pfg07) | GGCGGAGGTGTGCCGGAGGAATAGTGGCGGCGTGTGA | +1bp |
|  | 1 (Pfg19) | GGCGGAGTTGTGCCG-----TAGTGGCAGCGTGTGA | -5bp |
| #2-2 ($T_1$) | 1 (Pfg07) | GGCGGAGGTGTGCCGGAGGAATAGTGGCGGCGTGTGA | +1bp |
| | 2 (Pfg07) | GGCGGAGGTGTGCCG---GATAGTGGCGGCGTGTGA | -3bp |
|  | 1 (Pfg19) | GGCGGAGTT-----------TAGTGGCAGCGTGTGA | -11bp |
| #2-3 ($T_1$) | 1 (Pfg07) | GGCGGAGGTGTGCCGGAGGAATAGTGGCGGCGTGTGA | +1bp |
|  | 1 (Pfg19) | GGCGGAGTTGTGCCG-----TAGTGGCAGCGTGTGA | -5bp |
|  | 2 (Pfg19) | GGCGGAGTT-----------TAGTGGCAGCGTGTGA | -11bp |
| #2-4 ($T_1$) | 1 (Pfg07) | GGCGGAGGTGTGCCGGAGGAATAGTGGCGGCGTGTGA | +1bp |
| | 2 (Pfg07) | GGCGGAGGTGTGCCG---GATAGTGGCGGCGTGTGA | -3bp |
|  | 1 (Pfg19) | GGCGGAGTTGTGCCG-----TAGTGGCAGCGTGTGA | -5bp |
| #2-5 ($T_1$) | 1 (Pfg07) | GGCGGAGGTGTGCCGGAGGAATAGTGGCGGCGTGTGA | +1bp |
| | 2 (Pfg07) | GGCGGAGGTGTGCCG---GATAGTGGCGGCGTGTGA | -3bp |

| | | | | |
|---|---|---|---|---|
|  | 1 | (Pfg19) | GGCGGAGTTGTGCCG-----TAGTGGCAGCGTGTGA | -5bp |
| #2-6 ($T_1$) | 1 | (Pfg07) | GGCGGAGGTGTGCCGGAGGAATAGTGGCGGCGTGTGA | +1bp |
| | 2 | (Pfg07) | GGCGGAGGTGTGCCG---GATAGTGGCGGCGTGTGA | -3bp |
|  | 1 | (Pfg19) | GGCGGAGTTGTGCCG-----TAGTGGCAGCGTGTGA | -5bp |
| #3-1 ($T_1$) | 1 | (Pfg07) | GGCGGAGGTGTGCCGGAGGAATAGTGGCGGCGTGTGA | +1bp |
| | 2 | (Pfg07) | GGCGGAGGTGTGCCG---GATAGTGGCGGCGTGTGA | -3bp |
|  | 1 | (Pfg19) | GGCGGAGTTGTGCCG-----TAGTGGCAGCGTGTGA | -5bp |
|  | 2 | (Pfg19) | GGCGGAGTT-----------TAGTGGCAGCGTGTGA | -11bp |
| #3-2 ($T_1$) | 1 | (Pfg07) | GGCGGAGGTGTGCCGGAGGAATAGTGGCGGCGTGTGA | +1bp |
| | 2 | (Pfg07) | GGCGGAGGTGTGCCG---GATAGTGGCGGCGTGTGA | -3bp |
|  | 1 | (Pfg19) | GGCGGAGTTGTGCCG-----TAGTGGCAGCGTGTGA | -5bp |
| #3-3 ($T_1$) | 1 | (Pfg07) | GGCGGAGGTGTGCCGGAGGAATAGTGGCGGCGTGTGA | +1bp |
| | 2 | (Pfg07) | GGCGGAGGTGTGCCG---GATAGTGGCGGCGTGTGA | -3bp |
|  | 1 | (Pfg19) | GGCGGAGTTGTGCCG-----TAGTGGCAGCGTGTGA | -5bp |
| #3-4 ($T_1$) | 1 | (Pfg07) | GGCGGAGGTGTGCCGGAGGAATAGTGGCGGCGTGTGA | +1bp |
|  | 1 | (Pfg19) | GGCGGAGTTGTGCCG-----TAGTGGCAGCGTGTGA | -5bp |
|  | 2 | (Pfg19) | GGCGGAGTT-----------TAGTGGCAGCGTGTGA | -11bp |
| #3-5 ($T_1$) | 1 | (Pfg07) | GGCGGAGGTGTGCCGGAGGAATAGTGGCGGCGTGTGA | +1bp |
| | 2 | (Pfg07) | GGCGGAGGTGTGCCG---GATAGTGGCGGCGTGTGA | -3bp |
|  | 1 | (Pfg19) | GGCGGAGTTGTGCCG-----TAGTGGCAGCGTGTGA | -5bp |
| #3-6 ($T_1$) | 1 | (Pfg07) | GGCGGAGGTGTGCCGGAGGAATAGTGGCGGCGTGTGA | +1bp |
| | 2 | (Pfg07) | GGCGGAGGTGTGCCG---GATAGTGGCGGCGTGTGA | -3bp |
|  | 1 | (Pfg19) | GGCGGAGTTGTGCCG-----TAGTGGCAGCGTGTGA | -5bp |
| #3-7 ($T_1$) | 1 | (Pfg07) | GGCGGAGGTGTGCCGGAGGAATAGTGGCGGCGTGTGA | +1bp |
|  | 1 | (Pfg19) | GGCGGAGTTGTGCCG-----TAGTGGCAGCGTGTGA | -5bp |
| #3-8 ($T_1$) | 1 | (Pfg07) | GGCGGAGGTGTGCCGGAGGAATAGTGGCGGCGTGTGA | +1bp |
| | 2 | (Pfg07) | GGCGGAGGTGTGCCG---GATAGTGGCGGCGTGTGA | -3bp |
|  | 1 | (Pfg19) | GGCGGAGTTGTGCCG-----TAGTGGCAGCGTGTGA | -5bp |
|  | 2 | (Pfg19) | GGCGGAGTT-----------TAGTGGCAGCGTGTGA | -11bp |
| #3-9 ($T_1$) | 1 | (Pfg07) | GGCGGAGGTGTGCCG---GATAGTGGCGGCGTGTGA | -3bp |
|  | 1 | (Pfg19) | GGCGGAGTTGTGCCG-----TAGTGGCAGCGTGTGA | -5bp |
| #3-10 ($T_1$) | 1 | (Pfg07) | GGCGGAGGTGTGCCGGAGGAATAGTGGCGGCGTGTGA | +1bp |
| | 2 | (Pfg07) | GGCGGAGGTGTGCCG---GATAGTGGCGGCGTGTGA | -3bp |
|  | 1 | (Pfg19) | GGCGGAGTTGTGCCG-----TAGTGGCAGCGTGTGA | -5bp |
| #3-11 ($T_1$) | 1 | (Pfg07) | GGCGGAGGTGTGCCGGAGGAATAGTGGCGGCGTGTGA | +1bp |
| | 2 | (Pfg07) | GGCGGAGGTGTGCCG---GATAGTGGCGGCGTGTGA | -3bp |

| | | | | |
|---|---|---|---|---|
|  | 1 | （Pfg19） | GGCGGAGTTGTGCCG-----TAGTGGCAGCGTGTGA | -5bp |
|  | 2 | （Pfg19） | GGCGGAGTT-----------TAGTGGCAGCGTGTGA | -11bp |
| #3-12 ($T_1$) | 1 | （Pfg07） | GGCGGAGGTGTGCCG---GATAGTGGCGGCGTGTGA | -3bp |
|  | 1 | （Pfg19） | GGCGGAGTTGTGCCG-----TAGTGGCAGCGTGTGA | -5bp |
| #3-13 ($T_1$) | 1 | （Pfg07） | GGCGGAGGTGTGCCGGAGGAATAGTGGCGGCGTGTGA | +1bp |
| | 2 | （Pfg07） | GGCGGAGGTGTGCCG---GATAGTGGCGGCGTGTGA | -3bp |
|  | 1 | （Pfg19） | GGCGGAGTTGTGCCG-----TAGTGGCAGCGTGTGA | -5bp |
| #3-14 ($T_1$) | 1 | （Pfg07） | GGCGGAGGTGTGCCGGAGGAATAGTGGCGGCGTGTGA | +1bp |
| | 2 | （Pfg07） | GGCGGAGGTGTGCCG---GATAGTGGCGGCGTGTGA | -3bp |
|  | 1 | （Pfg19） | GGCGGAGTTGTGCCG-----TAGTGGCAGCGTGTGA | -5bp |
|  | 2 | （Pfg19） | GGCGGAGTT-----------TAGTGGCAGCGTGTGA | -11bp |
| #4-1 ($T_1$) | 1 | （Pfg07） | GGCGGAGGTGTGCCGGAGGAATAGTGGCGGCGTGTGA | +1bp |
| | 1 | （Pfg19） | GGCGGAGTTGTGCCGGAGGAATAGTGGCAGCGTGTGA | +1bp |
| #4-2 ($T_1$) | 1 | （Pfg07） | GGCGGAGGTGTGCCGGAGGAATAGTGGCGGCGTGTGA | +1bp |
| | 1 | （Pfg19） | GGCGGAGTTGTGCCGGAGGAATAGTGGCAGCGTGTGA | +1bp |
| #4-3 ($T_1$) | 1 | （Pfg07） | GGCGGAGGTGTGCCGGAGGAATAGTGGCGGCGTGTGA | +1bp |
| | 1 | （Pfg19） | GGCGGAGTTGTGCCGGAGGAATAGTGGCAGCGTGTGA | +1bp |
| #4-4 ($T_1$) | 1 | （Pfg07） | GGCGGAGGTGTGCCGGAGGAATAGTGGCGGCGTGTGA | +1bp |
| | 1 | （Pfg19） | GGCGGAGTTGTGCCGGAGGAATAGTGGCAGCGTGTGA | +1bp |
| #4-5 ($T_1$) | 1 | （Pfg07） | GGCGGAGGTGTGCCGGAGGAATAGTGGCGGCGTGTGA | +1bp |
| | 1 | （Pfg19） | GGCGGAGTTGTGCCGGAGGAATAGTGGCAGCGTGTGA | +1bp |
| #4-6 ($T_1$) | 1 | （Pfg07） | GGCGGAGGTGTGCCGGAGGAATAGTGGCGGCGTGTGA | +1bp |
| | 1 | （Pfg19） | GGCGGAGTTGTGCCGGAGGAATAGTGGCAGCGTGTGA | +1bp |
| #4-7 ($T_1$) | 1 | （Pfg07） | GGCGGAGGTGTGCCGGAGGAATAGTGGCGGCGTGTGA | +1bp |
| | 2 | （Pfg19） | GGCGGAGTTGTGCCGGAGGAATAGTGGCAGCGTGTGA | +1bp |

**Table S2. List of enzyme-coding genes involved in the representative anthocyanin biosynthetic pathway of *P. frutescens*.**

| Enzyme | Chromosome | Start | End | Strand | Related Transcript ID | TPM | | | |
|---|---|---|---|---|---|---|---|---|---|
| | | | | | | Wild Type | #2-1 ($T_1$) | #2-2 ($T_1$) | #2-3 ($T_1$) |
| PAL | Pfg00008 | 16070424 | 16075369 | + | Pfg00008_13430.1 | 6.0 | 7.0 | 7.6 | 9.3 |
| PAL | Pfg00014 | 9777787 | 9780760 | + | Pfg00014_09830.1 | 40.5 | 71.8 | 54.9 | 53.8 |
| PAL | Pfg00015 | 39355431 | 39358742 | - | Pfg00015_24630.1 | 133.7 | 222.0 | 177.0 | 139.0 |
| PAL | Pfg00016 | 16355222 | 16358906 | - | Pfg00016_12940.1 | 7.8 | 10.1 | 9.6 | 12.1 |
| PAL | Pfg00017 | 37266063 | 37269320 | - | Pfg00017_24040.1 | 155.3 | 259.7 | 220.4 | 151.9 |
| PAL | Pfg00018 | 9911770 | 9914816 | + | Pfg00018_10180.1 | 34.5 | 60.7 | 54.8 | 50.8 |
| C4H | Pfg00011 | 47332006 | 47334207 | + | Pfg00011_34950.1 | 30.0 | 67.2 | 56.1 | 37.0 |
| C4H | Pfg00011 | 51987028 | 51990622 | + | Pfg00011_39790.3 | 190.3 | 217.8 | 172.7 | 159.9 |
| C4H | Pfg00011 | 52007716 | 52009473 | + | Pfg00011_39800.1 | 0.0 | 0.1 | 0.0 | 0.0 |
| C4H | Pfg00012 | 5392339 | 5394076 | - | Pfg00012_06670.1 | 0.0 | 0.0 | 0.0 | 0.0 |
| C4H | Pfg00012 | 5410919 | 5414405 | - | Pfg00012_06680.1 | 218.2 | 261.7 | 221.0 | 221.8 |
| C4H | Pfg00012 | 9906059 | 9908321 | - | Pfg00012_11510.2 | 183.1 | 354.6 | 360.2 | 281.9 |
| C4H | Pfg00014 | 4436250 | 4440473 | + | Pfg00014_04760.1 | 0.0 | 0.0 | 0.0 | 0.0 |
| C4H | Pfg00018 | 4410721 | 4414072 | + | Pfg00018_05000.1 | 0.0 | 0.1 | 0.2 | 0.1 |
| 4CL | Pfg00001 | 16203275 | 16211922 | - | Pfg00001_13140.2 | 6.1 | 9.6 | 7.6 | 7.2 |
| 4CL | Pfg00001 | 55572404 | 55580679 | - | Pfg00001_36040.1 | 5.0 | 6.1 | 4.7 | 4.6 |
| 4CL | Pfg00001 | 73574582 | 73579373 | + | Pfg00001_54320.2 | 4.6 | 6.1 | 4.9 | 5.1 |
| 4CL | Pfg00003 | 42411460 | 42417999 | - | Pfg00003_19190.1 | 10.1 | 9.5 | 8.3 | 10.2 |
| 4CL | Pfg00005 | 22261017 | 22265835 | - | Pfg00005_11650.1 | 5.3 | 7.2 | 6.1 | 7.1 |

| | | | | | | | | | |
|---|---|---|---|---|---|---|---|---|---|
| 4CL | Pfg00009 | 26706261 | 26710512 | - | Pfg00009_14580.1 | 9.7 | 11.7 | 10.5 | 9.8 |
| 4CL | Pfg00010 | 26410170 | 26415796 | + | Pfg00010_12550.1 | 124.5 | 126.8 | 109.4 | 90.8 |
| CHS | Pfg00004 | 60934566 | 60935849 | + | Pfg00004_30460.1 | 0.0 | 0.0 | 0.0 | 0.0 |
| CHS | Pfg00004 | 60993957 | 60996280 | + | Pfg00004_30470.1 | 0.0 | 0.0 | 0.0 | 0.0 |
| CHS | Pfg00004 | 61014535 | 61017055 | + | Pfg00004_30480.1 | 0.0 | 0.4 | 0.0 | 0.0 |
| CHS | Pfg00005 | 8542945 | 8545283 | - | Pfg00005_06230.1 | 0.0 | 1.2 | 0.8 | 0.4 |
| CHS | Pfg00006 | 10563963 | 10566426 | - | Pfg00006_09450.1 | 0.1 | 0.0 | 0.1 | 0.0 |
| CHS | Pfg00006 | 10620424 | 10621628 | - | Pfg00006_09480.1 | 0.1 | 0.3 | 0.0 | 0.0 |
| CHS | Pfg00010 | 48795645 | 48797188 | + | Pfg00010_26770.1 | 0.0 | 0.2 | 0.1 | 0.1 |
| CHS | Pfg00010 | 48857726 | 49059278 | + | Pfg00010_26810.1<br>Pfg00010_26830.1 | 554.4 | 667.7 | 610.9 | 682.2 |
| CHS | Pfg00011 | 3619995 | 3622988 | - | Pfg00011_03580.1 | 6.8 | 11.5 | 13.3 | 6.0 |
| CHS | Pfg00012 | 58451544 | 58454966 | + | Pfg00012_44780.1 | 3.0 | 7.1 | 9.0 | 3.0 |
| CHS | Pfg00013 | 13362543 | 13364523 | - | Pfg00013_11960.1 | 1.3 | 1.8 | 2.5 | 1.4 |
| CHS | Pfg00013 | 13405349 | 13407496 | - | Pfg00013_11980.1 | 592.7 | 762.7 | 690.7 | 704.7 |
| CHS | Pfg00013 | 13527237 | 13529034 | - | Pfg00013_12040.1 | 6.8 | 15.6 | 13.1 | 9.4 |
| CHI | Pfg00002 | 22021888 | 22022825 | + | Pfg00002_12460.1 | 534.5 | 742.8 | 719.3 | 755.2 |
| CHI | Pfg00012 | 15599074 | 15600043 | - | Pfg00012_16170.1 | 647.4 | 1065.5 | 968.7 | 1017.4 |
| F3H | Pfg00007 | 8546568 | 8550589 | - | Pfg00007_05480.1 | 163.0 | 53.3 | 104.9 | 51.4 |
| F3H | Pfg00019 | 42719499 | 42723597 | + | Pfg00019_27030.1 | 78.7 | 27.0 | 18.0 | 16.2 |
| F3'H | Pfg00003 | 17342158 | 17345640 | - | Pfg00003_11050.2 | 99.2 | 193.5 | 158.1 | 165.9 |
| F3'H | Pfg00009 | 12974052 | 12976957 | + | Pfg00009_08970.1 | 105.4 | 183.3 | 150.2 | 168.6 |
| F3'H | Pfg00014 | 19441033 | 19445341 | + | Pfg00014_17090.1 | 2.3 | 1.9 | 1.9 | 1.5 |

| | | | | | | | | | |
|---|---|---|---|---|---|---|---|---|---|
| F3'H | Pfg00018 | 18844942 | 18849528 | - | Pfg00018_16930.1 | 5.6 | 11.2 | 9.7 | 7.4 |
| FSII | Pfg00011 | 45357386 | 45360075 | - | Pfg00011_33150.1 | 72.1 | 124.2 | 105.5 | 108.5 |
| FSII | Pfg00012 | 11776514 | 11778416 | + | Pfg00012_13140.1 | 12.5 | 17.8 | 26.1 | 15.4 |
| FSII | Pfg00012 | 11804012 | 11806846 | + | Pfg00012_13160.1 | 75.4 | 121.8 | 104.2 | 128.5 |
| FSII | Pfg00012 | 55250227 | 55251836 | - | Pfg00012_42340.1 | 0.0 | 0.0 | 0.0 | 0.0 |
| FSII | Pfg00012 | 56247055 | 56248666 | + | Pfg00012_43210.1 | 1.0 | 0.7 | 1.4 | 1.0 |
| FLS | Pfg00003 | 73807947 | 73809978 | + | Pfg00003_36060.1 | 0.8 | 1.8 | 1.0 | 1.1 |
| FLS | Pfg00009 | 60912737 | 60914160 | + | Pfg00009_33350.1 | 1.0 | 1.2 | 2.6 | 1.5 |
| DFR | Pfg00002 | 77196106 | 77198238 | + | Pfg00002_66420.3 | 313.2 | 491.4 | 332.3 | 446.2 |
| ANS | Pfg00014 | 15138302 | 15140452 | - | Pfg00014_13790.1 | 454.2 | 668.2 | 487.3 | 603.1 |
| ANS | Pfg00018 | 15209883 | 15211949 | - | Pfg00018_14070.1 | 311.6 | 533.8 | 403.1 | 435.9 |
| 3GT | Pfg00001 | 15945368 | 15946959 | + | Pfg00001_12990.1 | 53.6 | 97.5 | 72.2 | 84.0 |
| 3GT | Pfg00001 | 55235896 | 55237458 | + | Pfg00001_35760.1 | 110.7 | 151.1 | 128.1 | 139.5 |
| OMT3 | Pfg00010 | 3847608 | 3850298 | + | Pfg00010_03010.1 | 0.8 | 0.8 | 1.3 | 2.0 |
| OMT3 | Pfg00011 | 32351997 | 32354624 | - | Pfg00011_22370.1 | 15.2 | 20.8 | 18.2 | 16.6 |
| OMT3 | Pfg00012 | 25275029 | 25277633 | + | Pfg00012_24350.1 | 17.0 | 23.9 | 21.6 | 19.0 |
| OMT3 | Pfg00013 | 35135622 | 35137213 | + | Pfg00013_25930.1 | 9.4 | 16.7 | 21.4 | 14.7 |
| OMT3 | Pfg00020 | 2372825 | 2374480 | + | Pfg00020_01050.3 | 5.8 | 7.4 | 10.5 | 5.9 |
| Myb113b | Pfg00008 | 4238969 | 4241651 | + | Pfg00008_05210.1 | 0.3 | 1.6 | 1.4 | 0.3 |
| Myb-p1 | pFG00008 | 4399483 | 4403525 | - | Pfg00008_05380.1 | 19.3 | 49.6 | 34.3 | 37.6 |
| Myb-p1 | pFG00016 | 5027597 | 5028981 | - | Pfg00016_05140.1 | 37.9 | 136.3 | 106.8 | 73.6 |

**Table S3. Segregation of transgene in $T_1$ progeny.**

| $T_1$ progeny | Number of individuals | Transgene + | Transgene - | $\chi^2$-value | p-value | Fit to 3:1 ratio ($\alpha$=0.05) |
|---|---|---|---|---|---|---|
| #1 family | 49 | 38 | 11 | 0.17 | 0.68 | Yes |
| #2 family | 55 | 40 | 15 | 0.15 | 0.70 | Yes |
| #3 family | 49 | 38 | 11 | 0.17 | 0.68 | Yes |
| #4 family | 48 | 37 | 11 | 0.11 | 0.74 | Yes |

**Table S4. List of Oligonucleotides used in this study.**

| Purpose | Region | Oligonucleotides set | Sequence (5' to 3') | Fragment length (bp) | Fragment No. |
|---|---|---|---|---|---|
| Sanger sequencing | *F3H* | F3H_Seq1F | GAGTACCCTCCACCCCAAAT | 2349/2391 | — |
| | | F3H_Seq1R | CAGGCTTTGGAGAGACCATC | | |
| Construction of gRNA cassette | *F3H* | F3H_gRNA1F | ATTGGGAGTTGTGCCGGAGGATAG | — | — |
| | | F3H_gRNA1R | AAACCTATCCTCCGGCACAACTCC | | |
| Amplicon sequencing | *F3H* | F3H_Seq1F_tag | TCGTCGGCAGCGTCAGATGTGTATAAGAGACAG(NNNNNN)GAGTACCCTCCACCCCAAAT | 222/222 | — |
| | | F3H_Seq2R_tag | GTCTCGTGGGCTCGGAGATGTGTATAAGAGACAG(NNNNNN)GTCATCCCTTCAACGACCAT | | |
| Screening of transformation and null segregants | T-DNA | 00755F (F3H_gRNA1F) | ATTGGGAGTTGTGCCGGAGGATAG | 150 | |
| | | 00904R | GCGCTGTTATCAACCACTTTGTAC | | |
| Confirmation of null segregants | Non T-DNA | 00004F | CACGCCCTTTTAAATATCCGATTAT | 1496 | 1 |
| | | 01499R | ACCTCTAAAAACCTGTTCATTGTTA | | |
| | T-DNA | 00281F | GTAAAACGACGGCCAGT | 1219 | 2 |
| | | 01499R | ACCTCTAAAAACCTGTTCATTGTTA | | |
| | | 01197F | TCCAAATATCAACCCTGGCAA | 1713 | 3 |

| | | | | |
|---|---|---|---|---|
| | 02909R | CTTCTTCTGGCGGTTCTCTTCAGCC | | |
| | 02800F | CTGGGCAACACCGACCGGCACAGCA | 1500 | 4 |
| | 04299R | CATTCCCTCGGTCACGTATTTCACT | | |
| | 04200F | CCTGCCCAACGAGAAGGTGCTGCCC | 1501 | 5 |
| | 05719R | ACTCGCTTTCCAGCTTAGGGTACTT | | |
| | 05600F | AGGATTTCCAGTTTTACAAAGTGCG | 1520 | 6 |
| | 07116R | CGCGCCTTATCTTTAATCATATTCC | | |
| | 07012F | GTTTCTATAATCCATTGTGAATGTT | 1489 | 7 |
| | 08500R | GGGCACAACAGACAATCGGCTGCTC | | |
| | 08380F | GCCTCGTCCTGCAGTTCATTCAGGG | 671 | 8 |
| | 09050R | GTGATTTTGTGCCGAGCTGCCGGTC | | |
| Non T-DNA | 09026F | GACCGGCAGCTCGGCACAAAATCAC | 1449 | 9 |
| | 10474R | CCCAATTTGTGTAGGGCTTATTATG | | |
| | 14500F | TGGTGCGGTTTCATGCTTGTTCCTC | 987 | 10 |
| | 00004R | CACGCCCTTTTAAATATCCGATTAT | | |
| *F3H* | F3H_Seq1F | GAGTACCCTCCACCCCAAAT | 222/222 | 11 |
| | F3H_Seq2R | GTCATCCCTTCAACGACCAT | | |

**Table S5. HPLC conditions for the quantification of major compounds.**

| Compound | Extraction solvent | Standard compound (supplier, purity) | Column | Mobile phase (v/v) | Flow rate (mL/min) | Detection wavelength (nm) | Remarks |
|---|---|---|---|---|---|---|---|
| Perillaldehyde | 80% MeOH | Perillaldehyde（Nacalai Tesque Inc.,-) | Develosil ODS-HG (4.6 mm × 250 mm; Nomura) | MeOH : DW = 72 : 28 | 0.7 | 231 | – |
| Anthocyanins | 0.5% TFA | Cyanidin 3,5-diglucoside (ChromaDex Inc.,-) | Develosil ODS-HG (4.6 mm × 250 mm; Nomura) | A: AcOH : MeCN : DW : TFA = 8 : 10 : 81.5 : 0.5<br>B: AcOH : MeCN : DW : TFA = 20 : 25 : 54.5 : 0.5 | 1 | 520 | Linear gradient A → B over 30 min |
| Rosmarinic acid | 80% MeOH | Rosmarinic acid (Fujifilm Wako Pure Chemical Corp., ≥96% HPLC grade) | Unison UK-C18 (4.6 mm × 150 mm; Intakt) | 2.5% AcOH : MeCN : MeOH = 35 : 7 : 5 | 1 | 330 | – |
| Luteolin and its analogs | 80% MeOH | Luteolin (Fujifilm Wako Pure Chemical Corp., ≥95% Biochemical grade) | Unison UK-C18 (4.6 mm × 150 mm; Intakt) | 5% Formic acid : MeOH = 6 : 4 | 1 | 360 | – |

**Table S6. Analytical conditions for luteolin quantification using LC-QTOF/MS.**

| LC | |
|---|---|
| Model Name | 1260 Infinity II (Agilent Technologies, USA) |
| Mass Spectrometer | X500R QTOF (SCIEX, USA) |
| Column | ZORBAX Eclipse Plus C18 (2.1 mm × 100 mm, 3.5 μm; Agilent Technologies) |
| Column Temperature | 40 °C |
| Mobile Phase A | 0.1% formic acid in water |
| Mobile Phase B | 0.1% formic acid in methanol |
| Gradient B (%) | 0 min: 10% ; 3 min: 10% ; 18 min: 95% ; 25 min: 95% ; 25.1 min: 10% |
| Flow Rate | 0.2 mL/min |
| Injection Volume | 5 μL |
| **MS** | |
| Model Name | X500R QTOF (SCIEX, USA) |
| Ionization Mode | Electrospray ionization (ESI), negative |
| Monitoring Modes | TOF MS and TOF MS/MS (MRM HR) |
| Ion Spray Voltage | −4500 V |
| Source Temperature | 400 °C |
| Gas Settings | GS1: 60 psi; GS2: 50 psi; Curtain gas: 25 psi |
| Collision Energy | TOF MS: −10 V; TOF MS/MS: −35 V |
| Identification Criteria | Retention time and TOF MS/MS spectra |
| Quantification Method | Internal standard method using peak areas of luteolin (m/z 285.040) and isotope-labeled PFOS (m/z 506.957) |
| Calibration Range | 1–50 ng/mL (six-point calibration curve) |
