## Supplemental Figures for "Genome editing of the flavanone 3-hydroxylase gene produces a green *Perilla* line with elevated luteolin accumulation"

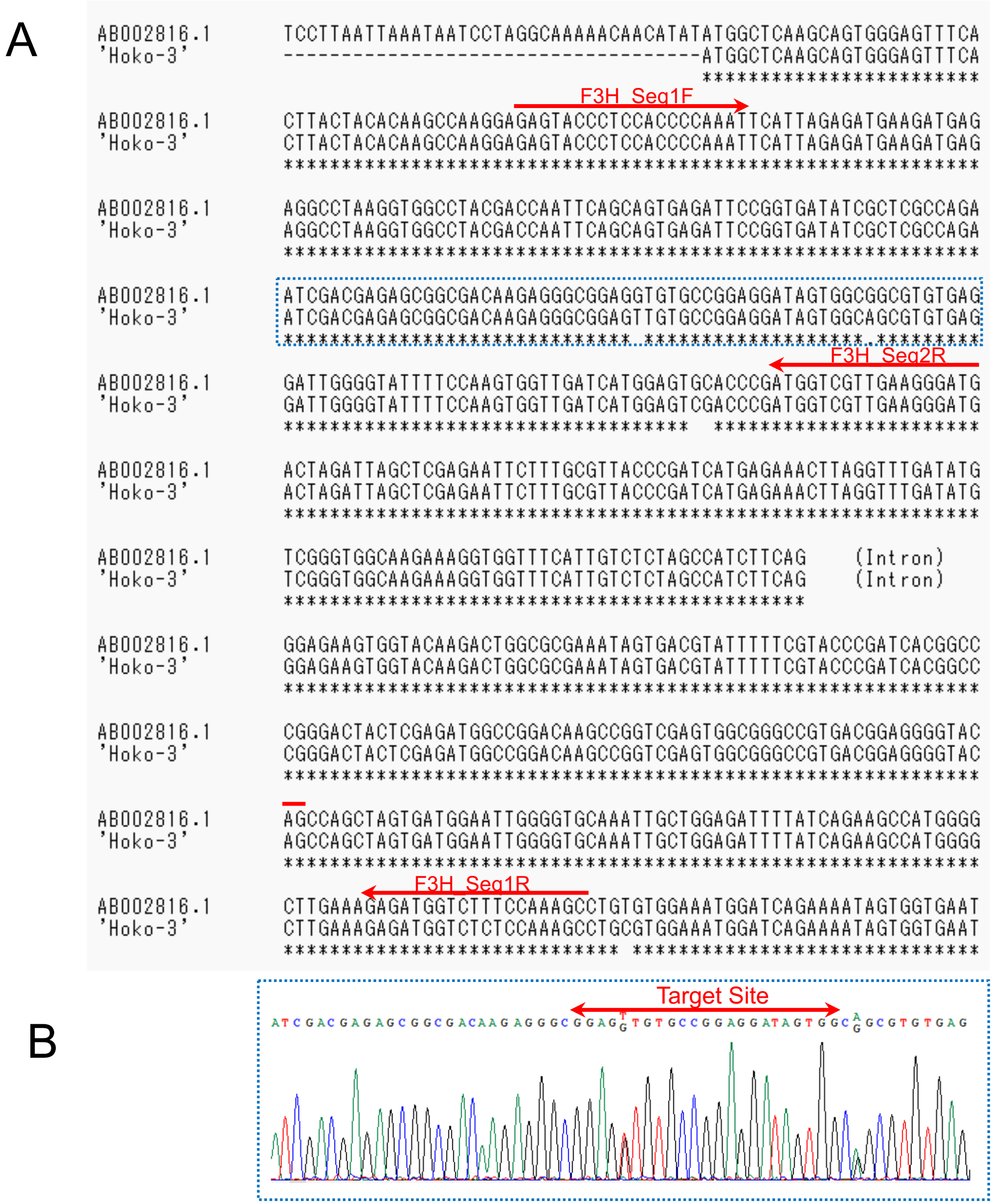


**Fig. S 1 Sequence organization of the Perilla F3H gene for gRNA design**

(A) Alignment of the CDS of the Perilla F3H gene with the 'Hoko-3' sequence reconstructed from RNA-seq data. Red arrows indicate the primers used for sequencing.

(B) Sanger sequencing of 'Hoko-3' in the region highlighted by the blue dashed box in panel (A). Red arrows indicate the target site and PAM sequence.

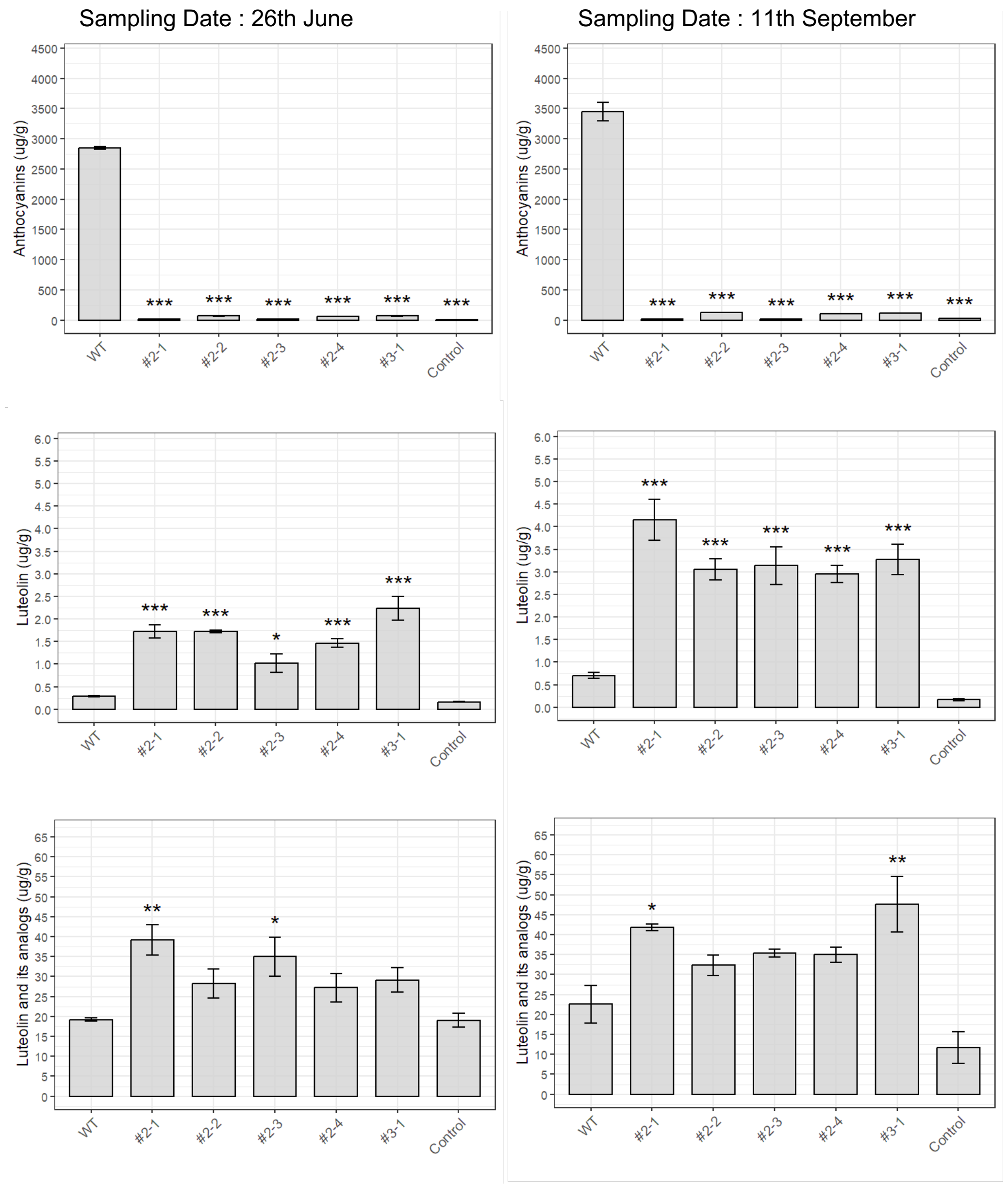


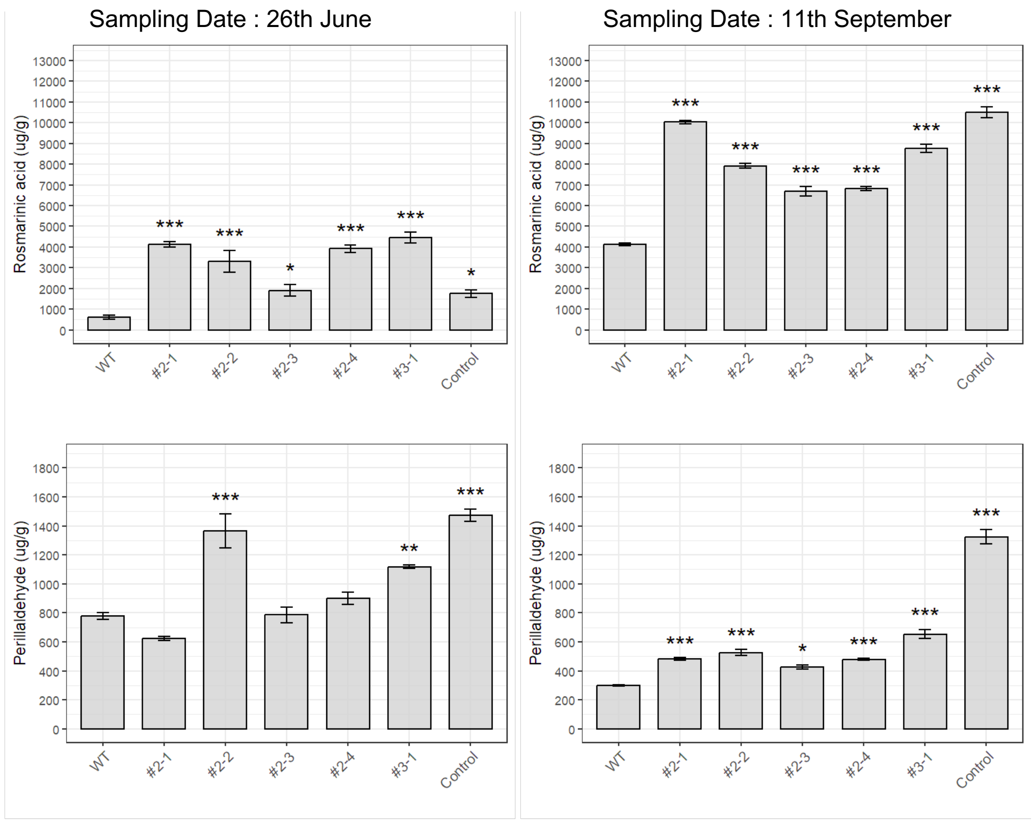


**Fig. S2. Metabolite accumulation in *Perilla frutescens* collected on June 26 (left panels) and September 11 (right panels).**
Quantification of five major metabolites—**Anthocyanins, Luteolin, Luteolin and its analogs, Rosmarinic acid, and Perillaldehyde**—in wild-type *P. frutescens* 'Hoko-3' (WT, red perilla), genome-edited lines (**#2-1, #2-2, #2-3, #2-4, #3-1**), and the non-red cultivar 'Ooba Ao-shiso' (**Control**, green perilla).
Bars indicate the mean ± SE of three biological replicates (n = 3).
Statistical analyses were performed separately for each compound and sampling date using one-way ANOVA followed by **Dunnett's test**, with WT ('Hoko-3') used as the reference group.
Asterisks denote significant differences relative to WT (*P < 0.05; **P < 0.01; ***P < 0.001).
